# A reproductive arrest program triggered by defects in Piwi and germ granules

**DOI:** 10.1101/276782

**Authors:** Maya Spichal, Bree Heestand, Katherine Kretovich Billmyre, Stephen Frenk, Shawn Ahmed

## Abstract

In several species, Piwi/piRNA genome silencing defects lead to immediate sterility accompanied by heterochromatin dysfunction and transposon-induced genomic instability, which may cause Piwi mutant sterility. In *C. elegans,* Piwi pathway mutants transmit a heritable stress through germ cells that induces sterility after growth for several generations. We found that sterile Piwi pathway mutant germ cells displayed inconsistent increases in DNA damage but consistently altered perinuclear germ granules that are known to promote fertility. Germ granule dysfunction did not elicit transposon expression but was sufficient to induce multiple phenotypes found in sterile Piwi silencing mutants, including germline atrophy and regrowth. Furthermore, loss of the germ granule component PGL-1 accelerated sterility in response to deficiency for *prg-1*/Piwi. Restoration of germ granule function to sterile *pgl-1* mutants restored their fertility. Together, our results suggest that germ granule defects may promote an adult reproductive arrest phenotype that is responsible for Piwi/piRNA mutant sterility.

## Introduction

Germ cells give rise to the mortal somatic tissues while maintaining themselves in a pristine condition that allows them to be propagated for an indefinite number of generations. This capacity for self-renewal is termed “germ cell immortality”, and functions to protect the germline from forms of damage that can result in sterility (Smelick and Ahmed, 2005a). Defects in *Caenorhabditis elegans* (*C. elegans)* genes that promote germline immortality cause a Mortal Germline (Mrt) phenotype, where robust fertility occurs for several generations but then deteriorates and finally culminates in complete sterility. The transgenerational sterility of *mrt* mutants implies that their germ cells may age as they proliferate over the generations such that they accumulate a form of damage that ultimately evokes sterility. Identifying the transgenerational damage that accumulates in *mrt* mutant germ cells and the mechanism(s) by which this damage induces sterility are central questions relevant to understanding the basis of germ cell immortality.

The initial germline immortality pathway defined in *C. elegans* was promoted by telomerase, a reverse transcriptase that maintains telomere length and stability by adding *de novo* telomere repeats to chromosome termini. *C. elegans* mutants that are defective for telomerase-mediated telomere repeat addition display progressive telomere erosion. In later generations, some telomeres become critically short and fuse together, leading to accumulation of end-to-end chromosome fusions (Lowden et al., 2008; Meier et al., 2006). The precise cause of telomerase mutant sterility remains uncertain (Karlseder et al., 1999; Nakamura et al., 1998; Sharpless and DePinho, 2007; Shay and Wright, 2005), but might be the creation of circular chromosomes that are toxic in meiosis (Nakamura et al., 1998).

Most human somatic cells can proliferate *in vitro* for a limited number of divisions before they enter a state of irreversible cell cycle arrest termed senescence (Sharpless and DePinho, 2007; Shay and Wright, 2005). Senescent cells are large and flattened in comparison to proliferating cells, they express senescence-associated beta-galactosidase and the tumor suppressor protein p16, and they secrete growth factors such as IL6 and IL8 cytokines that act in a cell-non-autonomous manner to promote aging (Tchkonia et al., 2013).The limited proliferative capacity of primary human somatic cells is called ‘replicative aging’. Lack of telomerase is thought to cause replicative aging in humans, because 1) most primary human somatic cells lack detectable telomerase activity *in vitro* (Counter et al., 1992; Shay and Wright, 2000), 2) primary human somatic cells experience progressive telomere erosion as they divide (Harley et al., 1990), 3) senescent cells display DNA damage response foci at telomeres that activate p53 to induce senescence (Fagagna et al., 2003; Herbig et al., 2004), and 4) because senescence does not occur if telomerase activity is restored by expression of the telomerase reverse transcriptase TERT, which is typically silenced in human somatic cells (Bodnar et al., 1998). Support for the model that telomere attrition induces senescence in humans was provided by acute knockdown of the telomere capping protein TRF2, which results in deprotection of telomeres that are normal in length, accompanied by DNA damage response foci at telomeres and senescence (Karlseder et al., 1999). Together, the above data support a model that telomere uncapping results in DNA damage response that activates p53 to cause senescence in human somatic cells.

Other forms of stress can accumulate to cause replicative aging of mammalian somatic cells. For example, expression of the p16 protein induces senescence and increases by orders of magnitude in mouse and human tissues with age, even though mouse somatic cells express telomerase (Krishnamurthy et al., 2004, 2006; Sharpless and DePinho, 2007). Forms of stress that accumulate during replicative aging of mammalian cells may be revealed by analysis of *C. elegans mortal germline* mutants, which display transgenerational sterility phenotypes that could reflect ‘replicative aging’ of germ cells. In this regard, deficiency for Piwi/piRNAs and related small RNA-mediated genome silencing proteins like RSD-6 and HRDE-1 results in transgenerational sterility in the absence of end-to-end chromosome fusions, indicating that telomere uncapping does not cause sterility of Piwi/piRNA genome silencing mutants (Billmyre et al., 2019; Buckley et al., 2012; Sakaguchi et al., 2014; Simon et al., 2014a).

P-M Hybrid Dysgenesis was discovered in the 1970’s as a temperature-sensitive *Drosophila* sterility defect that was linked to high levels of transposition (Kelleher, 2016; Kidwell et al., 1977), which occurs in response to lack of piRNA-mediated genome silencing of specific transposable elements (Brennecke et al., 2008a). Piwi is a conserved Argonaute protein that interacts with thousands of piRNAs in germ cells to repress expression of transposons and foreign nucleic acids that represent parasitic threats to the integrity of the genome (Aravin et al., 2007; Luteijn and Ketting, 2013; Siomi et al., 2011). Deficiency for Piwi proteins leads to immediate sterility in *Drosophila*, zebrafish and mouse, which correlates with increased transposon expression and DNA damage in sterile animals (Juliano et al., 2011; Kelleher, 2016; Siomi et al., 2011). Although this implies that transposon-induced genome instability could cause sterility of Piwi mutants, analysis of *Drosophila* strains with variable levels of transposons suggested that transposon-induced genome instability may only explain a minor fraction of P-M hybrid sterility (Srivastav and Kelleher, 2017). Therefore, the cause of sterility in Piwi mutants remains an unsolved problem in experimental biology.

Deficiency for the *C. elegans* Piwi ortholog *prg-1* compromises germ cell immortality (Simon et al., 2014b), rather than inducing the immediate sterility phenotype that is observed in Piwi mutants in some other species. The reason for the transgenerational delay in the onset of sterility in *prg-1*/Piwi mutants and in related Piwi pathway mutants of *C. elegans* is not clear, but it is possible that genomic silencing marks that are induced by the Piwi/piRNA pathway are more robustly maintained in germ cells of *C. elegans* if the Piwi silencing system is inactivated. The transgenerational damage that accumulates in *C. elegans* Piwi mutants may therefore correspond to progressive deterioration of Piwi-mediated heterochromatin.

Heterochromatin dysfunction in *C. elegans* has been associated with increased levels of transposon and tandem repeat expression, with RNA-DNA hybrid formation, and with transposon-induced DNA damage (McMurchy et al., 2017; Padeken et al., 2019; Zeller et al., 2016). However, very low levels of transposition occur in late-generation *prg-1* mutants, and *C. elegans mutator* mutants, which display high levels of transposition, do not become sterile if grown at low temperatures, suggesting that transposition is unlikley to cause transgenerational sterility of *prg-1*/Piwi mutants (Simon et al., 2014b).

Sterile late-generation *C. elegans prg-1*/Piwi mutants display an unusual pleiotropic germ cell degeneration phenotype as L4 larvae mature into 1 day old adults, such that most animals display ‘atrophied’ germlines that contain a small population of mitotic germ cells, whereas some germlines are ‘empty’ and lack germ cells altogether, some germlines are ‘short’ such that they have fewer germ cells than normal but possess both mitotic and meiotic germ cells, and some germlines are sterile but ‘normal’ in size such that they are not smaller than germlines of fertile late-generation *prg-1* mutant siblings (Heestand et al., 2018). Similarly, growth of *C. elegans* Piwi pathway mutants *rsd-6*, *nrde-2* and *hrde-1* at the restrictive temperature of 25°C for several generations results in sterile late-generation L4 larvae with germlines that are predominantly normal in size, but these larvae mature into sterile adults that possess germline arms with variable sizes, including small, atrophied and empty (Billmyre et al., 2019; Sakaguchi et al., 2014). The histone 3 lysine 4 demethylase RBR-2 also promotes germ cell immortality at 25°C and may promote genome silencing downstream of small RNAs (Alvares et al., 2014).

Atrophied germ lines of sterile *prg-1* mutants can regrow on day 2 of adulthood, and fertility can be restored to a minor fraction of sterile late-generation *prg-1* mutants by altering their food source (Heestand et al., 2018; Simon et al., 2014a). This implies that Piwi mutant sterility is a reversible form of reproductive arrest, also known as Adult Reproductive Diapause or Reproductive Quiescence, which can be induced in response to severe environmental stresses such as starvation (Angelo and Van Gilst, 2009; Padilla and Ladage, 2012; Tatar and Yin, 2001). We therefore proposed that late-generation sterility of *prg-1*/Piwi mutants may occur in response to accumulation of a heritable stress that is transmitted by germ cells (Heestand et al., 2018).

Some Argonaute proteins that regulate silencing of the epigenome, like PRG- 1/Piwi, are located in germ granules, which are perinuclear structures that interact with nuclear pore complexes and have been previously demonstrated to be necessary for fertility (Fig. 6A) (Updike et al., 2011). *C. elegans* germ granules are termed P granules based on antibodies that label the P blastomere that generates the germ cell lineage (Strome and Wood, 1982). Here we study the late-generation sterility phenotype of Piwi pathway genome silencing mutants and find that it is associated with abnormalities of germ granules. We show that germ granule dysfunction can induce reproductive arrest and that germ granule dysfunction interacts with Piwi deficiency to promote sterility.

## Results

### Genome instability in Piwi pathway genome silencing mutants

Transgenerational sterility in *C. elegans* telomerase mutants that lack the ability to maintain sequences that cap chromosome ends is caused by high levels of telomere fusions and associated genome damage (Meier et al., 2006). Many *mrt* mutants only become sterile when grown at the non-permissive temperature 25°C (Ahmed and Hodgkin, 2000), some of which are deficient for Piwi pathway-mediated genome silencing (Ashe et al., 2012; Buckley et al., 2012; Burkhart et al., 2011; McMurchy et al., 2017; Sakaguchi et al., 2014; Smelick and Ahmed, 2005b; Spracklin et al., 2017). However, the deficiency for the Piwi Argonaute protein PRG-1 or the nuclear RNA interference (RNAi) proteins NRDE-1 or NRDE-4 results in progressive sterility at any temperature (Batista et al., 2008; Buckley et al., 2012; Burkhart et al., 2011; Das et al., 2008; McMurchy et al., 2017; Simon et al., 2014b). We asked if DNA damage is directly associated with the sterility of Piwi pathway mutants by comparing fertile late-generation Piwi pathway mutants with late generation siblings that were sterile, using both temperature-sensitive Piwi pathway mutants that become sterile at 25°C and non-conditional mutants that become sterile when grown at 20°C.

Although DNA damage has been suggested not to contribute significantly to the sterility of *prg-1* mutants (Simon et al., 2014b), DNA damage signaling in sterile *prg-1* mutant animals has not been previously studied. We therefore examined early and late-generation *prg-1* mutants grown at 20°C across several generations for activation of the DNA damage response, as detected by an antibody to phosphorylated S/TQ, a protein modification that is created by two sensors of DNA damage, ATM and ATR, which are phosphatadylisonisol-3-kinase like protein kinases (Kim et al., 1999; Vermezovic et al., 2012). We observed a significant increase in the fraction of germ cells with upregulated DNA damage response for late-generation fertile and sterile *prg-1* mutants (∼3- to 6- fold) when compared to either wild type controls or to early-generation *prg-1* mutants (Fig. 1A,B). However, the fraction of germ cells with an activated DNA damage response was not consistently elevated in sterile *prg-1*/Piwi mutants in comparison to fertile late-generation siblings (Fig. 1B). We also tested temperature-sensitive Piwi pathway mutants and found that the wild-type DNA damage response was upregulated by growth at 25°C but that this level was not further elevated in germlines of sterile late-generation *rsd-6, nrde-2* or *hrde-1* mutant adults (Fig. 1B,C).

**Figure 1:**
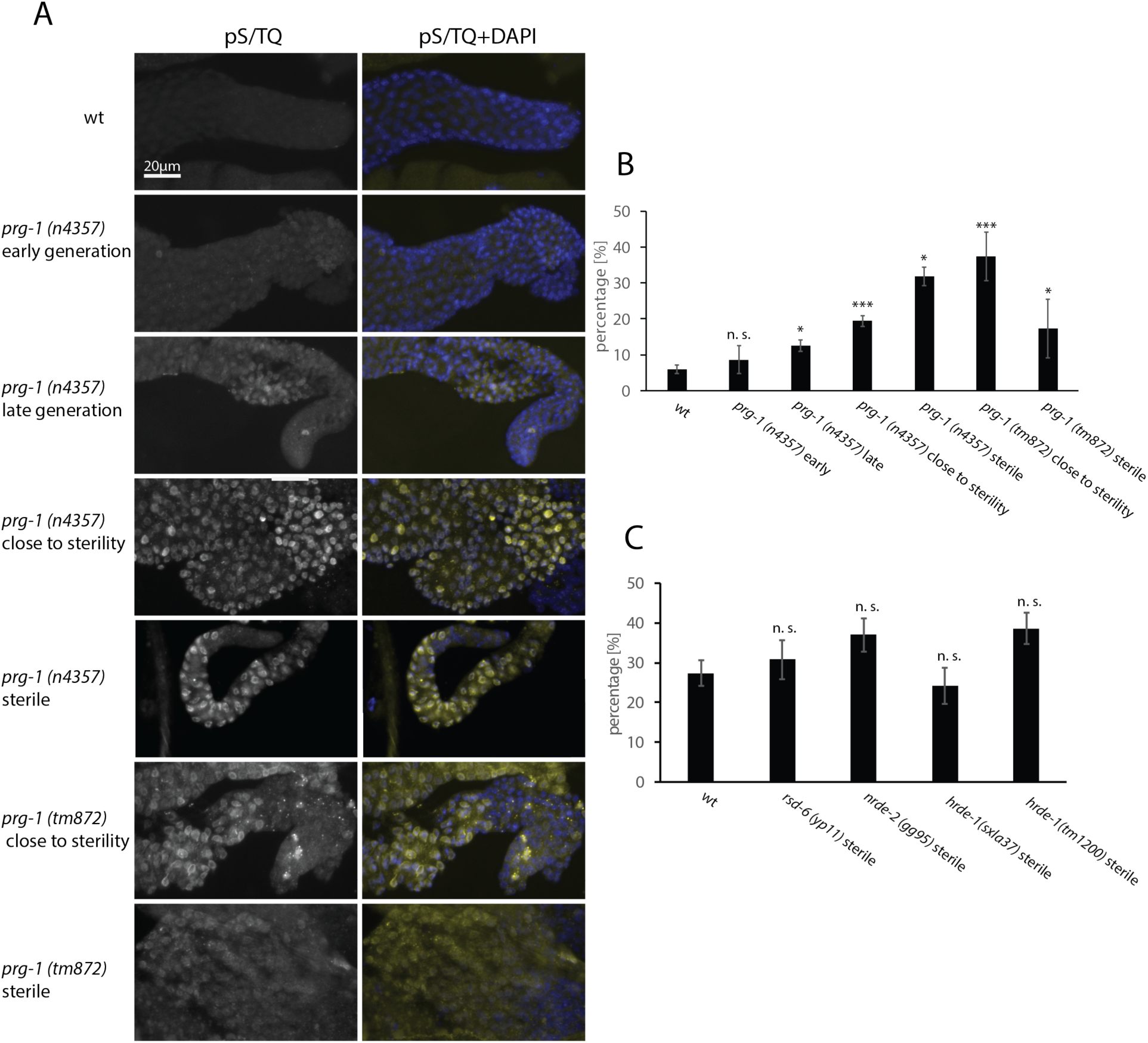
DNA Damage Response is variable in Piwi pathway genome silencing mutants prior to and at sterility. **(A)** Staining of pS/TQ in wild type, *prg-1(n4357) and prg-1(tm876)* mutants. pS/TQ(grey and yellow) and DAPI(blue) **(B)** Quantification of pS/TQ staining in 20°C wild type and *prg-1* mutants at different levels of fertility. **(C)** Quantification of pS/TQ staining in 25°C wild type and small RNA genomic silencing mutants at sterility. P-values were obtained by comparing sample groups in a Mann-Whitney U test (p-value: P > 0.05 = n. s., P < 0.05 = *, P = <0.01 = **, P =< 0.001 = *** compared to wild type).

### Piwi pathway genome silencing mutants exhibit germ granule defects at sterility

If secondary siRNAs are depleted from *prg-1* mutants and then restored by crossing distinct *prg-1; mutator* double mutants together, this causes immediate sterility for *prg-1* mutant F1 cross progeny that possess germ cells with germ granule defects and a range of adult germline sizes that are similar to those of sterile late-generation Piwi pathway mutants (de Albuquerque et al., 2015; Phillips et al., 2015). Furthermore, combined loss of the epigenomic regulators SPR-5 and LET-418, or loss of H3K9 methylation modifier SET-2 and the nuclear RNAi pathway, results in immediate sterility that is associated with disruption of germ granules (P granules in *C. elegans*) (Käser-Pébernard et al., 2014; Robert et al., 2014). Although most Piwi pathway mutants do not display severe fertility defects until the generation of sterility, late-generation *prg-1*/Piwi mutants display very small brood sizes for many generations prior to sterility (Heestand et al., 2018). By immunofluorescence with a P granule antibody, we observed that P granules were frequently strongly reduced or absent in nuclei of sterile late-generation *prg-1* mutant germ cells (hereafter referred to as marked P granule dysfunction) (Fig. 2B, S1, Table 1), which was less common in fertile *prg-1(n4357)* mutant siblings while *prg-1(tm872)* mutants showed more consistent P granule defects right before and at sterility (Fig. 2B-B’, Table 1). Consistently, growth of the temperature-sensitive Mrt mutants *rsd-6*, *nrde-2*, *hrde-1* and *rbr-2* at the restrictive temperature of 25°C resulted in sterile late-generation animals with marked P granule dysfunction, whereas their fertile late-generation siblings did not exhibit pronounced P granule defects (Fig. 2C, D, E, C’, D’, E’, S1, Table 1).

**Figure 2:**
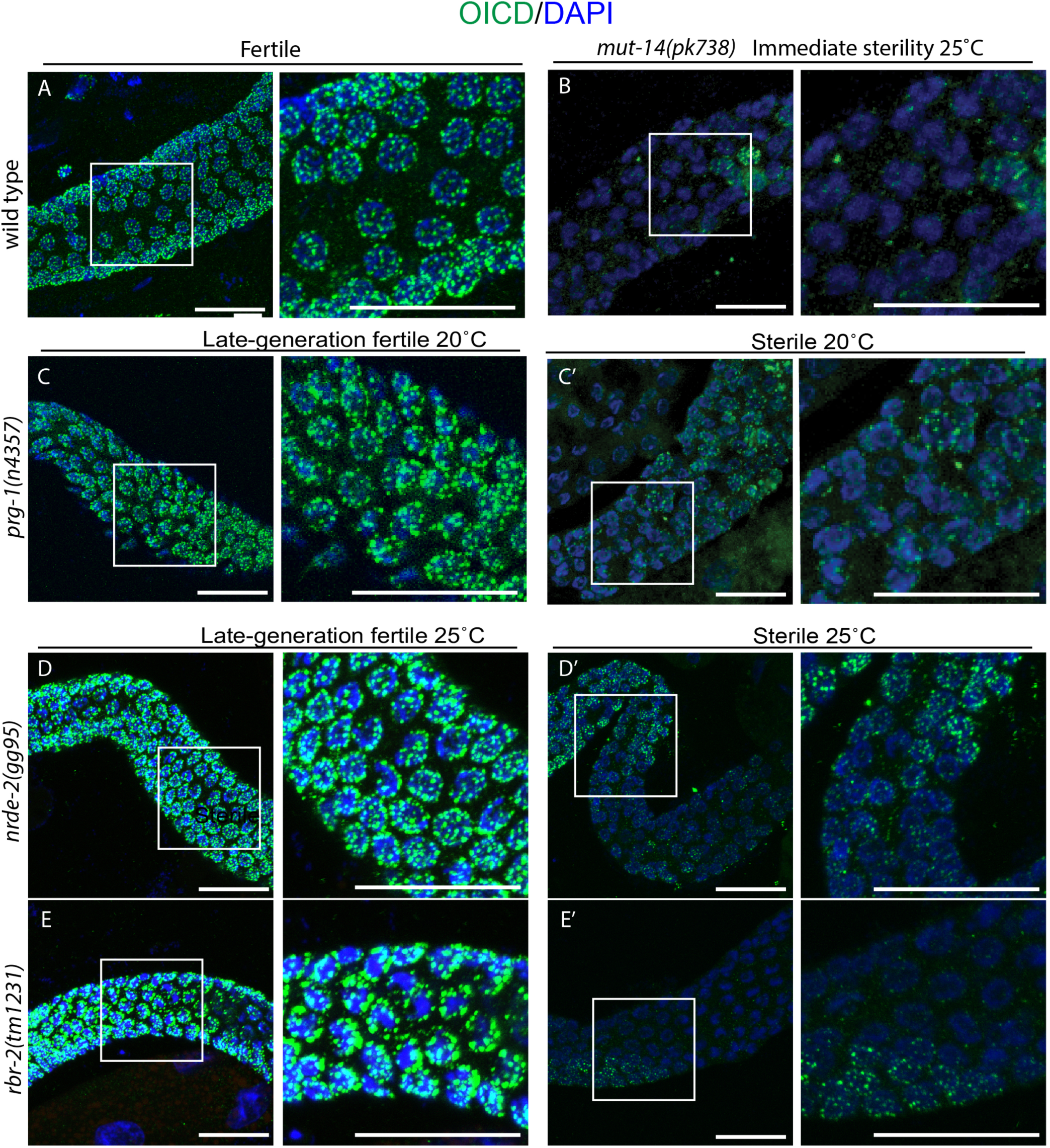
At sterility, partial loss of P granule staining occurred in *prg-1* and temperature-sensitive mutants. Germlines of sterile Day 2-3 adult animals were stained using the OIC1D4 antibody against P granules (green) and DAPI (blue). **(A)** Control animals contain uniform puncta of P granule staining surrounding each nuclei.(B) *mut-14* mutants display an immediate sterility when shifted to 25°C for one generation and display P granule abnormalities **(C-E)** Late generation fertile *prg-1,nrde-2* and *rbr-2* animals displayed relatively normal P granule staining in the germlines. **(C’- E’)** *prg-1*, *nrde-2 and rbr-2* sterile animals all exhibited some degree of germ cell P granule loss.

**Table 1:** Quantification of P granule defects in the germline. Sterile Piwi pathway genome silencing mutants were stained with P granule antibody. Germlines arms were semi-qualitatively scored for P granules. If 20% or more of the the germline arm exhibited P granules abnormalities (complete or partial loss of P granules) germlines were scored as abnormal.

|  | fertile | sterile |
| --- | --- | --- |
|  | 20°C |  |
| wt (n=12) | 0.00% | NA |
| *prg-1(n4357)* (n=16) | 36.36% | 100.00% |
| *prg-1(tm872)* (n=21) | 87.50% | 92.31% |
| *nrde-4(gg131)* (n=13) | 50.00% | 90.91% |
| *nrde-1(yp4)* (n=13) | 20.00% | 87.50% |
|  | 25°C |  |
| wt (n=35) | 0.00% | NA |
| *mut-14(pk738)*<br>(n=15) | NA | 85% |
| *rsd-6(yp11)* (n=32) | 0.00% | 40.00% |
| *nrde-2(gg95)* (n=25) | 0.00% | 100.00% |
| *rbr-2(tm1231)* (n=32) | 0.00% | 100.00% |
| *hrde-1(tm1200)*<br>(n=31) | 0.00% | 100.00% |

Germ line atrophy occurs for many but not all sterile Piwi pathway mutants as L4 larvae develop into adults (Billmyre et al., 2019; Heestand et al., 2018). Large-scale apoptosis contributes to but is not essential for this germ cell degeneration phenotype (Heestand et al., 2018), and P granules have been shown to vanish in cells that undergo apoptosis (Min et al., 2016; Sung et al., 2017). However, we observed marked P granule dysfunction in germlines of sterile late-generation *nrde-2* and some *rsd-6* mutant adults that were wild type in size and had not undergone significant amounts of apoptosis (Fig. 2D, D’; Fig. S1F, F’). *nrde-1* and *nrde-4* mutant animals displayed a sterility phenotype where many sterile germline arms displayed P granule defects (90% and 87% respectively) (Fig. S1, Table 1). P granule defects in *nrde* mutants included P granule dissociation from the nuclear envelope into the cytoplasm and P granule aggregations as well as P granule loss. Therefore, dysfunction of *nrde-1* and *nrde-4*, which act downstream of multiple small RNA genome silencing pathways (Fig. 6A), might influence P granules differently than we observed in the germline arms of most sterile Piwi pathway mutants.

*C. elegans* P granules are adjacent to Mutator bodies, which house Mutator proteins that recruit RNA-dependent RNA polymerase to create secondary effector siRNA populations in response to primary siRNAs (Phillips et al., 2012). *mutator* mutants are dysfunctional for secondary siRNA biogenesis and remain fertile indefinitely at low temperatures (Simon et al., 2014b), but display an immediate highly penetrant fertility defect at 25°C that could be caused by dysfunction of Piwi pathway-mediated heterochromatin (Ketting et al., 1999). We examined germlines of sterile *mut-14(pk738)* mutant adults at the restrictive temperature of 25°C and observed pronounced P granule defects in 66.67% of animals with no offspring and 100% P granule defects in animals that laid eggs that could not develop into adults, here referred to as ‘fertile’(Fig. 2B’, Table 1).

### Germ granule dysfunction phenocopies sterile Piwi pathway genome silencing mutants

We addressed the significance of P granule dysfunction in sterile late-generation small RNA genome silencing mutants using an RNA interference clone that simultaneously targets four P granule subunits *pgl-1*, *pgl-3*, *glh-1*, *glh-4* (Fig. 6A) (Knutson et al., 2017; Updike et al., 2014). This resulted in many second generation (G2) sterile adults at 25°C (80%) (Fig. 3A). The germlines of G2 P granule RNAi L4 larvae were normal in size in comparison to control worms. However, a pronounced germ cell atrophy phenotype occurred as many P granule-depleted animals matured into sterile adults, resulting in a significant change in their germline profile (Fig. 3B, p=3.93E-24).

**Figure 3:**
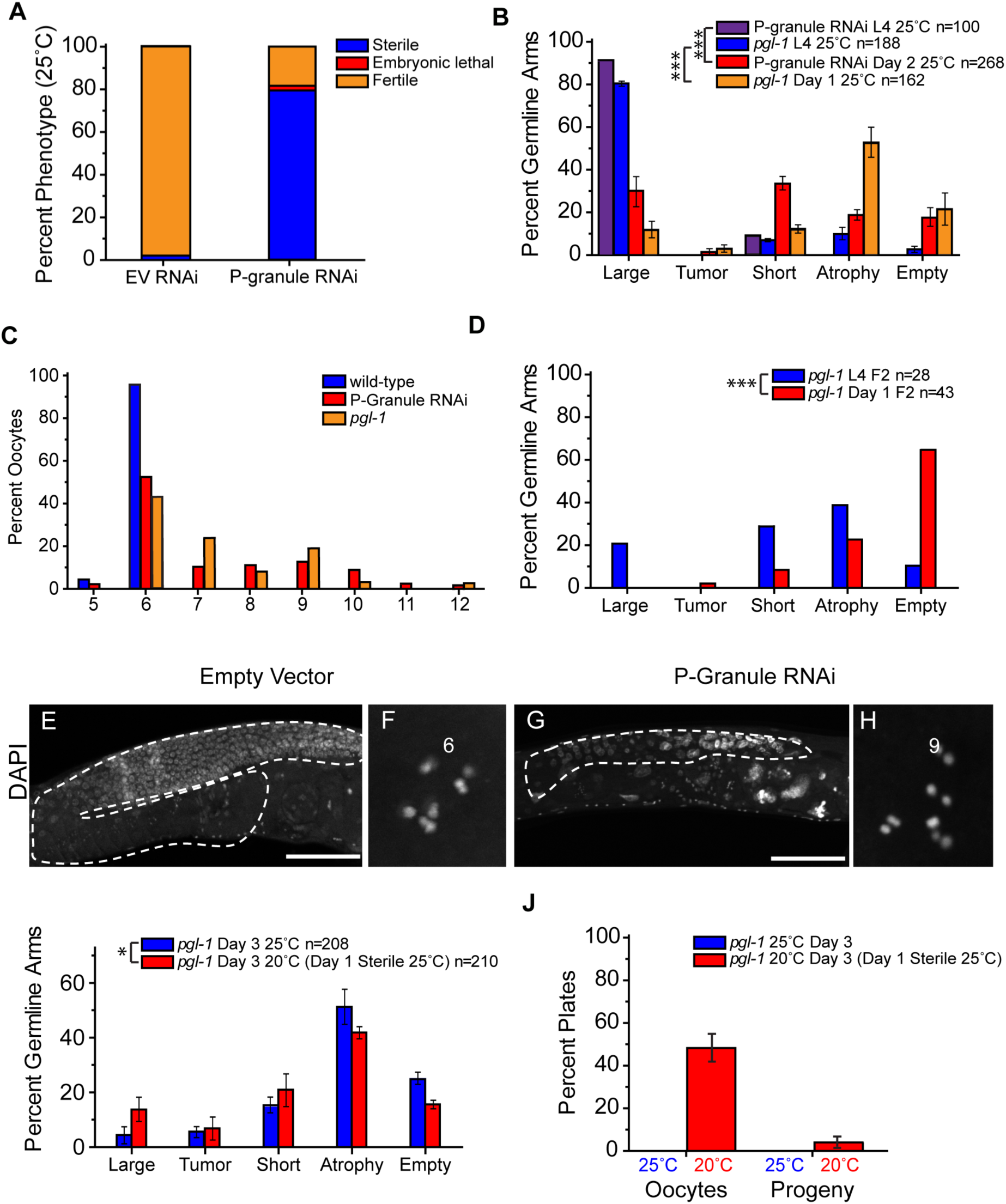
Disruption of P granules by RNAi or by deficiency for *pgl-1* results in similar phenotypes. **(A)** G2 wildtype (N2) worms on Empty Vector (EV) or P granule RNAi at 25°C were scored for sterility, embryonic lethality, or fertility. n=16-30 worms per condition, 2 independent experiments. **(B)** Germline phenotypes of G1 L4 *pgl-1* mutants and G2 L4 P granule RNAi treated animals stained by DAPI. **(C)** G1 *pgl-1* mutants and G2 P granule RNAi treated animals contained oocytes with increased numbers of univalents at sterility as measured by DAPI bodies n=151 wild type, 81 P granule RNAi, 37 *pgl-1*. **(D)** Stained DAPI germlines of sterile G2 L4 and Day 1 *pgl-1* mutants. **(E-H)** Representative DAPI images of germline and oocyte defects in empty vector treated animals **(E,F)** and P granule RNAi treated animals **(G,H)** scale bar=10µm. **(I)** *pgl-1* G1 worms at 25°C were either DAPI stained as Day 3 adults (blue) or shifted to 20°C at Day 1 and DAPI stained at Day 3 (red). **(I)** No sterile day 1 *pgl-1* G1 at 25°C laid oocytes or progeny (n=35 plates, 350 worms). Sterile day 1 *pgl-1* shifted to 20°C (n=50 plates, 500 worms) gave rise to plates with oocytes and rarely progeny. Plates with oocytes contained both oocytes and dead embryos. Plates with progeny contained embryos that hatched to L1. For B,D,I *** p<0.0001, * p=0.007 Fisher’s Chi Square test with Bonferroni correction, n=total germline arms scored, error bars are S.E.M.

Furthermore, oocyte univalents were previously observed in sterile but not fertile late-generation *rsd-6* mutants (Sakaguchi et al., 2014), and 45.5% of oocytes of sterile P granule depleted adults contained 7-12 univalents (Fig. 3C, E-H), consistent with a previous report (Billmyre et al., 2019).

The germline phenotypes observed in P granule-depleted worms mimicked those of sterile *prg-1*/Piwi mutant adults (Fig. 3A-C), which can regrow their atrophied germlines and become fertile (Heestand et al., 2018). We tested the hypothesis that germ granule dysfunction might be sufficient to cause reproductive arrest using a mutant deficient for the P granule component *pgl-1*, which remains fertile at low temperatures but displays a pronounced sterility phenotype if shifted to the restrictive temperature of 25°C (Kawasaki et al., 1998). Most first generation (G1) *pgl-1* animals at 25°C displayed normal germlines at the L4 larval stage, but there was a significant shift in the germline profiles with 52% of day 1 adults showing germline atrophy (Fig. 3B, p=1.04E-35), similar to P granule RNAi knockdown and sterile *prg-1* mutants (Heestand et al., 2018). Furthermore, when oocytes of sterile *pgl-1* mutant adults were scored, we found that 56.7% of oocytes had univalents, in agreement with the oocyte univalents observed in response to P granule depletion by RNAi (Fig. 3C).

Some G1 progeny of *pgl-1* mutants whose parents were shifted from 20°C to 25°C were fertile and gave rise to G2 25°C animals that were uniformly sterile. 25°C G2 *pgl-1* mutant L4 larvae had germline profiles that displayed more severe germ cell proliferation defects than those of G1 L4 larvae, and day 1 adults showed a striking 54% increase in empty germlines (Fig. 3D, p=2.44E-05). Therefore, germlines of G1 progeny of *pgl-1* mutant mothers that were shifted to the restrictive temperature of 25°C for a single generation mimicked the germ cell degeneration phenotypes of sterile Piwi pathway mutants as L4 larvae matured into adults, whereas 25°C G2 generation *pgl-1* mutant larvae displayed more pronounced germ cell proliferation defects.

We asked if the fertility of sterile 25°C G1 *pgl-1* mutants could be restored by shifting them to the permissive temperature of 20°C. Sterile G1 *pgl-1* mutants that lacked embryos were shifted to 20°C on day 1 of adulthood and their germlines were scored after 48 hours, which revealed a significant change in their germline profiles (p=0.007), with a shift from atrophied germlines towards large germlines when compared to sterile G1 *pgl-1* sibling controls that remained sterile at 25°C (Fig. 3I). In agreement with the regrowth of some germline arms, we observed that some sterile G1 *pgl-1* mutant adults that were shifted to 20°C for several days had laid oocytes and dead embryos, and that a few even gave rise to progeny (Fig. 3J). In contrast, sterile *pgl-1* mutant control adults that were kept at 25°C never gave rise to oocytes, dead embryos or living progeny (Fig. 3J).

### Transcriptional consequences of germ granule dysfunction and small RNA genome silencing mutant sterility

The sterility of Piwi/piRNA mutants correlates with expression of transposons and associated genome instability (Brennecke et al., 2008b; Juliano et al., 2011; Kelleher, 2016; Siomi et al., 2011). We therefore studied RNA from the nuclear RNAi defective mutants *nrde-1*, *nrde-2* and *nrde-4* (Buckley et al., 2012; Guang et al., 2008, 2010) that have Mortal Germlines at 25°C (Burkhart et al., 2011). However, at 20°C *nrde-1* and *nrde-4* mutants become progressively sterile but *nrde-2* mutants remain fully fertile indefinitely (Burkhart et al., 2011). *nrde-1* and *nrde-4* mutants have relatively large brood sizes at sterility at 20°C, such that large cohorts of L4 larvae that are poised to become sterile can be collected. In contrast, *prg-1* mutants develop very low brood sizes in late generations (Heestand et al., 2018), which makes it difficult to obtain many synchronous animals that are poised to become sterile. We therefore prepared RNA from early-generation and sterile-generation *nrde-1* and *nrde-4* mutants at 20°C and compared this with RNA from wild type and *nrde-2* mutant controls.

We found that sterile generation *nrde-1* and *nrde-4* mutant L4 larvae showed strong upregulation of similar transposon classes in comparison to early generation *nrde-1* or *nrde-4* mutant L4 larvae or to wild type controls (Fig. 5A). We identified 18 transposon classes that were upregulated at least two-fold in both *nrde-1* and *nrde-4*, 13 of which were CER retrotransposons, which is consistent with previous transgenerational analysis of *hrde-1* mutants (Ni et al., 2016). However, we also found that *nrde-2* mutant controls that do not become sterile at 20°C displayed strong upregulation of similar transposon loci, even in early generations. Of the 18 transposons upregulated in both *nrde-1* and *nrde-4*, 17 were also upregulated in *nrde-2* compared with wild type. Previous work reported that distinct classes of transposons are upregulated in *prg-1* and *nrde-2* mutants (Das et al., 2008; McMurchy et al., 2017; Ni et al., 2016; Simon et al., 2014b). Consistently, we analyzed published RNA-seq data from *prg-1(n4357)* mutant animals (McMurchy et al., 2017) found that MIRAGE1 was the only transposon upregulated in both *nrde-1*, *nrde-4* and *prg-1* mutants (McMurchy et al., 2017) (Fig. 4A). Therefore, although deficiency for *prg-1* and *nrde-1/nrde-4* disrupts small RNA-mediated genome silencing and compromises germ cell immortality, these genes have distinct functions with regards to small RNA-mediated regulation of the genome.

**Figure 4:**
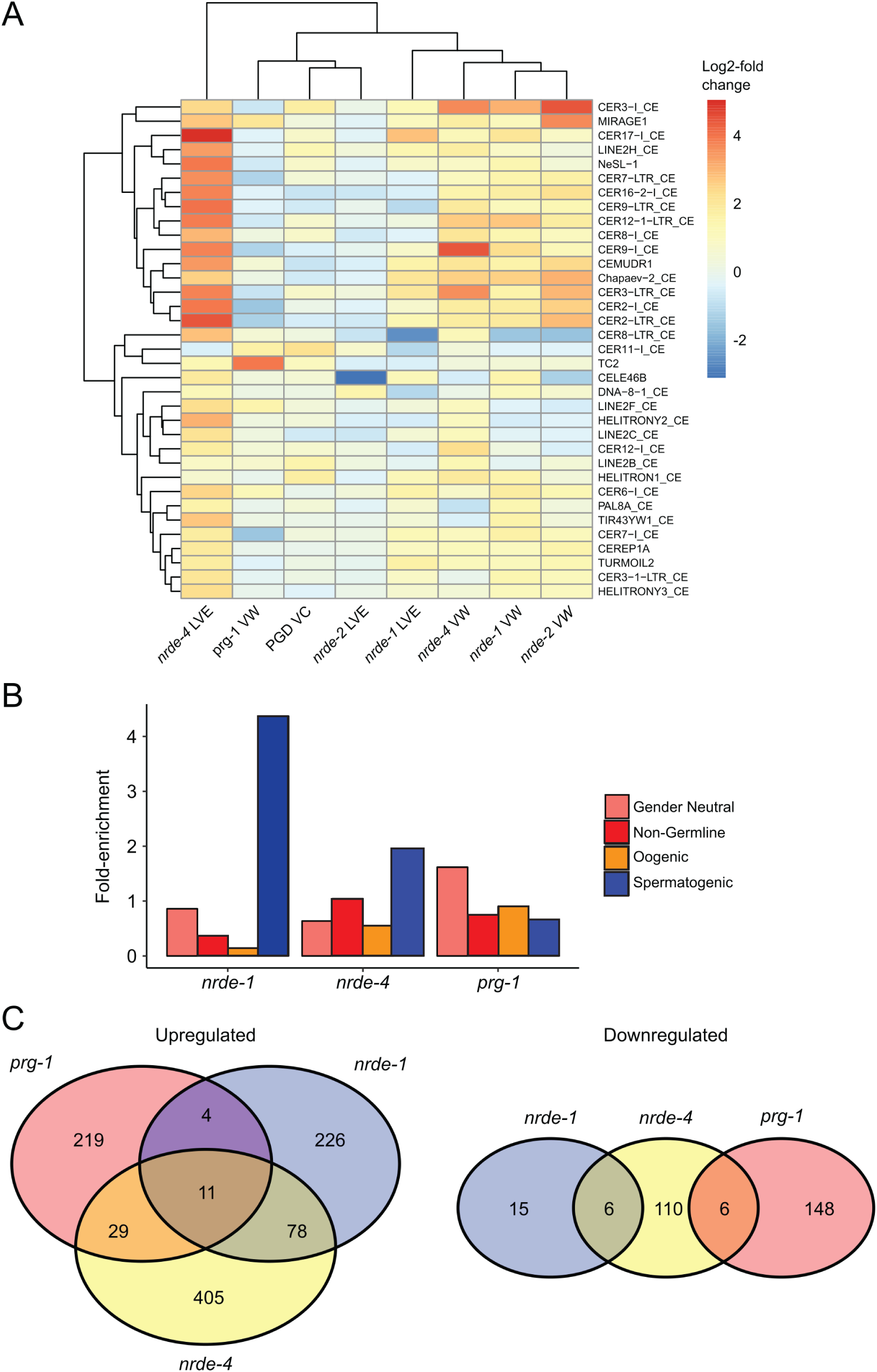
Transcriptome profiling of germ granule deficient larvae of Piwi pathway genome silencing mutants *nrde-1* and *nrde-4*. **(A)** Log2-fold changes in transposon transcript abundance in mutant vs wild type (VW), late vs early generation (LVE) or P granule RNAi-treated vs control (PGD VC). Transposons upregulated at least two-fold in *nrde-1*, *nrde-4* or *prg-1(n4357)* are shown. **(B)** Enrichment of germline-expressed genes in genes upregulated in *nrde-1* and *nrde-4* mutants. **(C)** Number of genes up and downregulated in *nrde-1* and *nrde-4* mutants.

**Figure 5:**
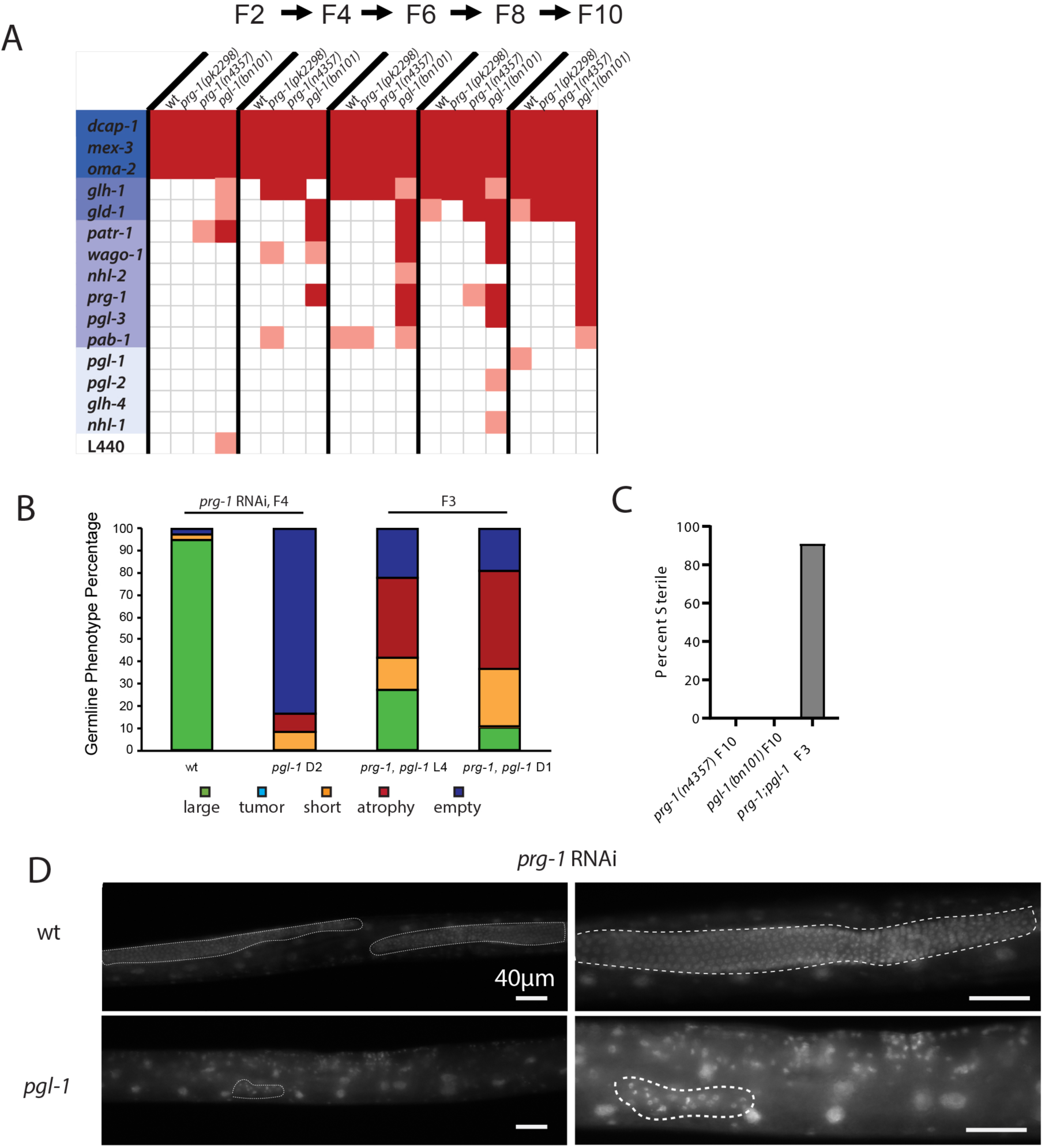
Deficiency for *pgl-1* results in rapid sterility of *prg-1* mutants. (A) Germline immortality assay with wild type, *prg-1(pk2298), prg-1(n4357)* and *pgl-1(bn101)* on different P-granule RNAi clones. 2 replicates of 3 L1 larvae were transferred every week or after 2 generations and plates were scored as sterile when no more than 3 worms were present after 1 week. Light red indicates sterility for 1 of 2 replicate plates while dark red indicates sterility for both replicates. When one plate became sterile worms from the second plate were used for 2 replicates. (B) Germline sizes of wild type and *pgl-1(bn101)* on *prg-1* RNAi after 4 generations and of *prg-1; pgl-1* double mutant F3 animals. (C) *prg-1; pgl-1* double mutants display marked sterility in the F3 generation.(D) Germline comparison of wild type and *pgl-1(bn101)* on *prg-1* RNAi after 4 generations.

We compared gene expression data from sterile generation *nrde-1* or *nrde-4* mutant L4 larvae with late-generation *prg-1* mutants (McMurchy et al., 2017) and with wild type or *nrde-2* mutant control L4 larvae in order to identify common genes whose expression is specifically induced in *nrde* mutant L4 larvae that are poised to become sterile and that might explain the P granule abnormalities at sterility. Late-generation *prg-1* mutant and sterile generation *nrde-1* and *nrde-4* mutant gene expression was very different from wild type or *nrde-2* mutant controls RNA. Spermatogenic genes were over-represented among genes significantly upregulated in sterile generation *nrde-1* and *nrde-4* mutant larvae (p = 1e-15, Chi Square test for both mutants), which is consistent with what has been previously reported for *spr-5* mutants that become progressively sterile (Katz et al., 2009). However, there was no general tendency of an up- or down-regulation of any gene category for *prg-1*, *nrde-1* and *nrde-4* mutants. Only 11 genes were significantly upregulated in all three mutants, and there were no commonly downregulated genes, which suggests that *prg-1* and *nrde-1/nrde-4* mutants have fundamentally different gene expression profiles at or near sterility (Fig. 4C). We identified a set of 20 genes that were significantly upregulated at least fourfold in *prg-1* and either *nrde-1* or *nrde-4*, which included *pud* (protein upregulated in dauer) genes that might be consistent with a role for DAF-16/Foxo in *prg-1* mutant sterility (Table S2) (Buckley et al., 2012; Simon et al., 2014a). However, we could not find consistent changes in any gene whose knockdown perturbs P granule structure (Updike and Strome, 2010), which suggests that altered germ granule structure of sterile Piwi pathway mutants is a post-transcriptional response.

RNA-seq data sets were previously obtained for germlines of P granule defective L4 larvae that were poised to become sterile, but gene expression was not significantly different from wild type (Knutson et al., 2017). We compared RNA from germlines of P granule defective L4 larvae (Knutson et al., 2017) with RNA from sterile generation *nrde-1* and *nrde-4* L4 larvae and found no genes that were up or downregulated in both *nrde-1/nrde-4* and P granule defective larvae. Moreover, only two of 18 transposons upregulated in *nrde-1/nrde-4* were also upregulated in P granule defective L4 larvae (Fig. 4A). We conclude although acute dysfunction of P granules based on simultaneous knockdown of four P granule components leads to germ line degeneration phenotypes that mimic those of sterile generation Piwi pathway mutants, it does not cause immediate overt effects on expression of heterochromatic segments of the genome or on genes in a manner that mimics the transcriptomes of Piwi pathway mutants that are poised to become sterile. Interestingly, the differences between *nrde-1/nrde-4* RNA data and *prg-1* also correlate with differences that we observed in P granule abnormalities between these mutants (Figure 2, S1). This indicates that the sterility observed in nrde and PIWI mutants causes P granule defects resulting from different molecular pathways.

### Deficiency for *pgl-1* enhances *prg-1*/Piwi mutant sterility

Given that deficiency for *pgl-1* or RNAi knockdown of P granule components at 25°C mimics Piwi mutant sterility, we performed an RNAi screen for P granule components whose knockdown induces sterility in early-generation *prg-1* mutants or in *pgl-1* mutants grown at the permissive temperature of 20°C. We knocked down several P-granule components for 10 generations in wild type control animals and in strains deficient for either *prg-1* or the P-granule component *pgl-1* (Fig 5A). Several RNAi clones led to immediate sterility for all genotypes tested, and we also identified P granule components whose knockdown induced sterility the *pgl-1* mutant. Specifically, we found that RNAi of *prg-1*/Piwi induced sterility in *pgl-1* mutants in generation F4 (Fig 5B). We studied the germline phenotypes of sterile *pgl-1* mutants treated with *prg-1* RNAi and observed that they had a mixture of empty, atrophied and short germlines (Fig. 5B, D).

To confirm the interaction of *pgl-1* mutation with RNAi knockdown of *prg-1*, we used outcrossed mutations in *prg-1* and *pgl-1* to create *prg-1; pgl-1* double mutants and compared them to the single mutant outcrossed controls. We found that *prg-1* (20/20) and *pgl-1* (20/20) single mutant controls were fertile for 10 generations when grown at 20°C. In contrast, *prg-1; pgl-1* double mutant F2 adults were fertile but gave rise to F3 animals that were almost all sterile (41/45) (Fig 5C). We stained sterile *prg-1; pgl-1* double mutant F3 with DAPI and found that on Day 1 62.9% of germlines were either atrophied or empty (Fig 5B).

## Discussion

Previous work has demonstrated that Piwi mutant sterility correlates with transposon expression, transposition and genomic instability in several species (Juliano et al., 2011; Kelleher, 2016; Siomi et al., 2011). We studied DNA damage signaling in *C. elegans* Piwi pathway mutants and found that this was not consistently increased in sterile animals in comparison to fertile late-generation siblings. These data complement previous observations that suggest that DNA damage induced by transposition is unlikely to cause the transgenerational sterility of *prg-1*/Piwi mutants (Simon et al., 2014b). A recent study revealed that defects in H3K9 methylation caused by *met-2* and *set-25* histone methyltransferases cause R-loops and mutations at tandem repeats in the *C. elegans* genome, reduced fertility, and synthetic sterility with the ATR homolog *atl-1* (Padeken et al., 2019) However, the sterility of the Piwi mutants we studied was not associated with high levels of ATR signaling, suggesting that Piwi sterility may be distinct from the reduced fertility caused by loss of H3K9me2 and me3 (Padeken et al., 2019).

We discovered pronounced germ granule defects in sterile but to a lesser extent in fertile late-generation Piwi pathway mutants and tested their significance by disrupting germ granule proteins (Fig. 3). We found that shifting a *pgl-1* germ granule mutant to restrictive temperature, led to germline degeneration phenotypes that phenocopied those of sterile generation Piwi pathway mutants (Heestand et al., 2018). We also found that restoring germ granule function sterile *pgl-1* mutant adults, by shifting them to the permissive temperature of 20°C, resulted in growth of the germline and fertility (Fig. 3I-J).

Although sterility in response to germ granule dysfunction has previously been considered to be a pathological condition that results in a germline-to-soma transformation (Knutson et al., 2017), our results indicate that germ granule dysfunction can induce reproductive arrest in adults (Fig. 3, 6C). As environmental stresses such as starvation can also evoke reproductive arrest (Angelo and Van Gilst, 2009; Padilla and Ladage, 2012), we speculate that germ granule dysfunction might represent a general mechanism for orchestrating stress-induced reproductive arrest.

Sterile late-generation *prg-1*/Piwi pathway mutants display a broad range of germline phenotypes that are observed in response to germ granule dysfunction, including germ cell atrophy, empty germlines, univalent chromosomes in oocytes, germline regrowth and reproductive arrest (Heestand et al., 2018). Together with the germ granule defects observed in sterile Piwi pathway mutants, our results suggest that Piwi mutant sterility is a form of reproductive arrest that may be caused by germ granule dysfunction (Fig. 6B). If the germ granule defects in sterile Piwi pathway mutants were a consequence rather than a cause of Piwi mutant sterility, then we would have expected that disrupting germ granule function would lead to a distinct form of sterility that did not bear hallmarks of Piwi mutant sterility like germ cell atrophy and reproductive arrest (Fig. 6C). Another reason that germ granule defects of sterile Piwi pathway mutants are unlikely to be a consequence of Piwi mutant sterility is that we observed some sterile Piwi pathway mutants with large germlines that did not display obvious P granule defects (Fig. 2, S1). We also found that deficiency for the P granule component *pgl-1* dramatically accelerated the sterility in response to deficiency for *prg-1*/Piwi, indicating that these proteins function redundantly to promote germ cell immortality. Knockdown of the germ granule components *gld-1* and *pgl-3* also induced F4 generation sterility for *pgl-1* mutants, which suggests a common function PRG-1, GLD-1 and PGL-3 in promoting transgenerational fertility in the context of germ granules that is consistent with their known physical interactions (Akay et al., 2013; Chen et al., 2016)

**Figure 6:**
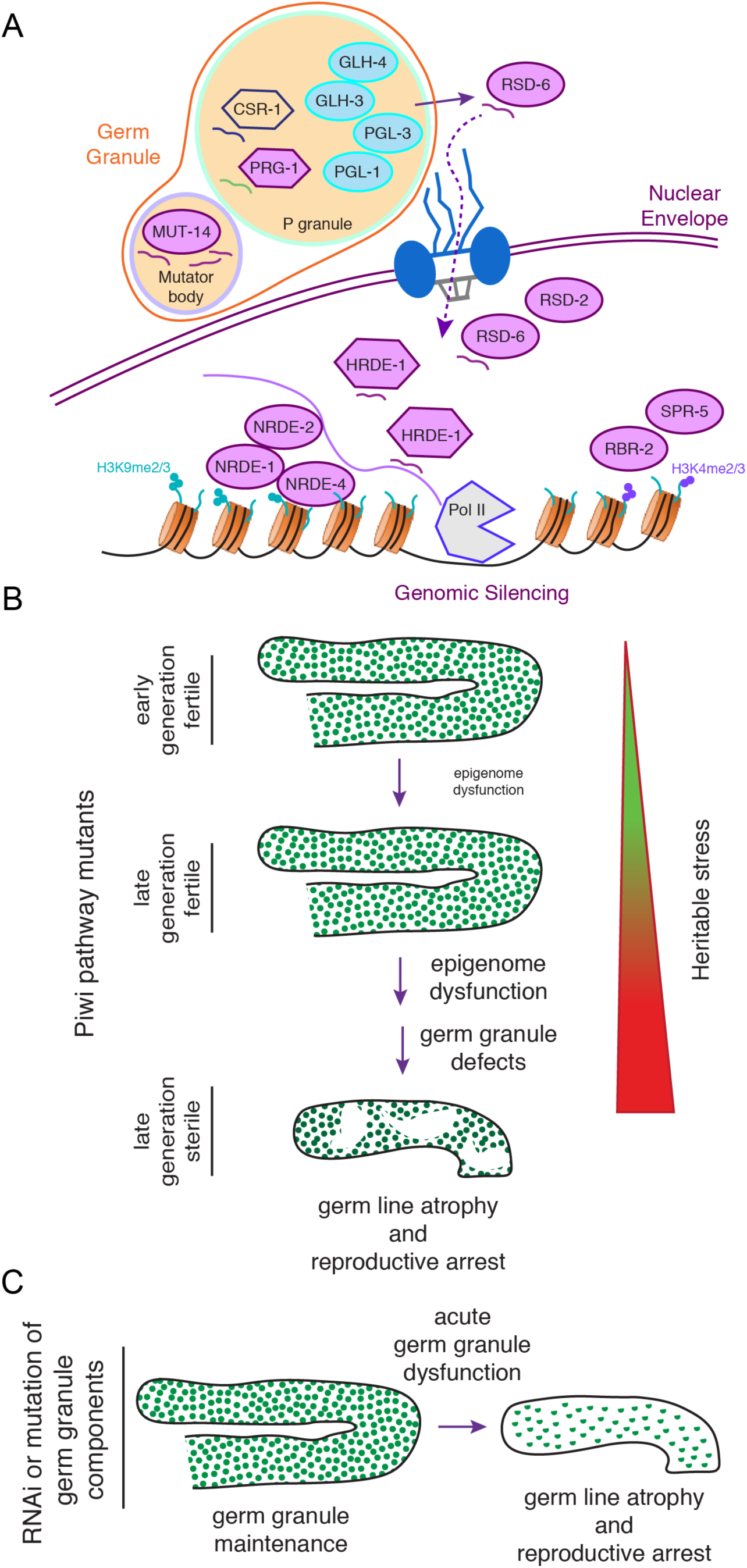
Model for P granule dysfunction in Piwi pathway genome silencing mutants. (**A**) Model of P granules and associated small RNA-mediated genome silencing proteins. Blue circles contain P granule components used in this paper. Mutants used this paper are in lavender. Argonaute proteins are hexagons. Non-conditional mutants have red outlines. **(B)** Model for transgenerational sterility in response to accumulation of Heritable Stress in Piwi pathway genome silencing mutants, which ultimately compromises P granule integrity. **(C)** Model for induction of reproductive arrest by acute dysfunction of P granule components.

Although P granules and associated Mutator bodies contain proteins like PRG- 1/Piwi and a host of other factors that promote small RNA biogenesis (Billi et al., 2012), we did not observe strong expression of heterochromatic transposons that are normally silenced by small RNAs in L4 larvae that are poised to become sterile because of germ granule dysfunction. This observation is consistent with the premise that small RNA-induced genomic silencing marks can be maintained in the absence of small RNA biogenesis.

Previous analysis showed that crosses that restored secondary siRNAs to *prg-1* mutants created sterile F1 *prg-1/*Piwi mutant progeny that possessed adult germlines with sizes analogous to those of sterile late-generation *prg-1* mutants and in disruption of P granules (de Albuquerque et al., 2015; Heestand et al., 2018). This led to the hypothesis that misrouting of small RNAs associated with pro- and anti-silencing small RNA pathways could be the cause of sterility in Piwi pathway genome silencing mutants (de Albuquerque et al., 2015; Phillips et al., 2015), which agrees with our observations of germ granule abnormalities in sterile but not fertile late-generation Piwi pathway mutants. Consistently, sterile mouse mutants with defects in piRNA biogenesis have been reported to display germ granule defects (Chuma et al., 2006; Zheng and Wang, 2012). We suggest that germ granule dysfunction could explain the sterility observed in P-M hybrid dysgenesis and the immediate sterility phenotype of Piwi mutants in several species (Juliano et al., 2011; Kelleher, 2016; Siomi et al., 2011).

When RNA from sterile generation *nrde-1* and *nrde-4* mutant L4 larvae and late-generation *prg-1*/Piwi mutants was examined, we failed to observe consistent depletion of any gene whose knockdown is known to disrupt P granule formation (Updike and Strome, 2010), suggesting a post-transcriptional reproductive arrest mechanism (Fig. 4B). We propose that transgenerational dysfunction of the Piwi/piRNA pathway in *C. elegans* results in heterochromatin defects and transcription of ‘stressful RNA’ from one or more genomic loci in later generations, which triggers a reproductive arrest program that perturbs germ granule structure (Fig. 6B). A role for defective heterochromatin as a cause of Piwi mutant sterility is supported by the *C. elegans* mutants *spr-5* and *rbr-2*, which encode H3K4 demethylases that function within nuclei downstream of small RNAs to promote genomic silencing (Fig. 2). *spr-5* and *rbr-2* mutants display transgenerational sterility phenotypes that mirror those of Piwi pathway silencing mutants (Alvares et al., 2014; Billmyre et al., 2019).

With regards to the model of telomerase deficiency as a cause of replicative aging in mammalian cells, we suggest that germ granule dysfunction in Piwi mutants may be analogous to telomere uncapping in the context of senescence (Sharpless and DePinho, 2007; Shay and Wright, 2000). Acute germ granule dysfunction is sufficient to mimic Piwi mutant sterility (Fig. 3, 6C), similar to knockdown of the TRF2 telomere capping protein being sufficient to induce acute telomere uncapping and senescence (Karlseder et al., 1999). In the case of replicative aging of mammalian cells, progressive telomere erosion is the source of damage that ultimately induces telomere uncapping. In the case of transgenerational Piwi pathway mutant sterility, it remains unclear what is the precise source of ‘heritable stress’ that accumulates over generations to induce sterility, although this is likely to be related to transgenerational desilencing of heterochromatin/transposons (Fig. 5A).

Our results establish a novel framework for considering inheritance in the context of epigenetic stress and how this can evoke sterility. We suggest that Piwi mutant sterility is a response to epigenome dysfunction that arrests germ cell development in an effort to protect the genome or epigenome in times of stress. Transposons and viruses are likely to be in a continuous arms race between parasite and host, where components of germ granules that promote Piwi pathway genome silencing and protect the host are likely to be targeted (Updike and Strome, 2010). It may therefore be reasonable to hypothesize the evolution of a reproductive arrest mechanism in the context of an attack on the Piwi silencing system that is predicated on disrupting the integrity of germ granules themselves.

## Materials and Methods

### Strains

All strains were cultured at 20°C or 25°C on Nematode Growth Medium (NGM) plates seeded with *E. coli* OP50. Strains used include Bristol N2 wild type, *hrde-1(tm1200) III, nrde-1(yp4) III, nrde-2(gg95) II, nrde-4 (gg131) IV, prg-1 (n4357) I, prg-1 (tm872) I, prg-1(pk2298) I, pgl-1(bn101) IV, mut-14 (pk738) V, rbr-2(tm1231) IV,* HIS-72::mNeonGreen (LP230).

### RNAi Assay

Feeding RNAi plates harboring host bacteria HT115(DE3) engineered to express *“*quad*”* dsRNA (targeting *pgl-1*, *pgl-3*, *glh-1*, *glh-4*) were obtained from Susan Strome (Knutson et al., 2017; Updike et al., 2014). L1 larvae were placed onto freshly prepared feeding RNAi plates with dsRNA induced by 1 mM IPTG (isopropyl-β-D(-)- thiogalactopyranoside) and were transferred after 1 generation at 25°C and collected at G2 adults as described in Knutson et al 2017 (Knutson et al., 2017) for DAPI staining, oocyte and germline analysis.

For RNAi screen, L1 worms containing the *his-72*:: *mNeonGreen* transgene were arrested in M9 media and then pipetted at a density of 20 worms per plate and maintained at 20°C. At the L4 stage and during adulthood worms were scored for germline size using Leica M205 fluorescence microscope.

For the P-granule germline immortality assay 3 L1 worms were transferred every week on two replicate plates per condition at 20°C. Plates were counted as sterile when no more than 3 worms were present after one week. Partial sterility was counted when one out of two plates became sterile while both replicate plates were sterile for full sterility.

### DAPI staining and Scoring

DAPI staining was performed as previously described (Ahmed and Hodgkin, 2000). Briefly, L4 larvae were selected from sibling plates and sterile adults were singled as late L4s, observed 24 hours later for confirmed sterility, and then stained 48 hours after collection. Univalents were scored by counting DAPI bodies in the −1 to −4 oocytes. Germline profiles were scored using the method outlined in Heestand et al 2018 (Heestand et al., 2018).

### Statistical Analysis

Statistical analysis was performed as previously described (Heestand et al., 2018). Briefly, statistical analysis was performed using the R statistical environment (R Core Team, 2013). For germline phenotypes, contingency tables were constructed and pairwise Chi Square tests with Bonferroni correction was used to determine significant differences in in germline phenotype distributions. The significance of the DNA damage response analysis was determined by a Kruskal-Wallis test, followed by a Mann-Whitney test between individual samples. P-values were adjusted with Bonferroni correction when multiple comparisons were performed.

### RNA extraction and sequencing

Animals were grown at 20°C on 60 mm NGM plates seeded with OP50 bacteria. RNA was extracted using Trizol (Ambion) followed by isopropanol precipitation. Library preparation and sequencing was performed at the UNC School of Medicine High-Throughput Sequencing Facility (HTSF). Libraries were prepared from ribosome-depleted RNA and sequenced on an Illumina Hiseq 2500.

### RNA-seq Analysis

The following publicly available RNA-seq datasets were download from the Gene Expression Omnibus (https://www.ncbi.nlm.nih.gov/geo/): GSE92690 (P granule RNAi experiment) and GSE87524 (*prg-1* experiment). Adapter trimming was performed as required using the bbduk.sh script from the BBmap suite (Bushnell) and custom scripts. Reads were then mapped to the *C. elegans* genome (WS251) using hisat2 (Kim et al., 2013) with default settings and read counts were assigned to protein-coding genes using the featureCounts utility from the Subread package (Liao et al., 2014). For multimapping reads, each mapping locus was assigned a count of 1/n where n=number of hits. Differentially expressed genes were identified using DESeq2, and were defined as changing at least 2-fold with FDR-corrected p-value < 0.01. For analysis of transposon RNAs, reads were mapped to the *C. elegans* transposon consensus sequences downloaded from Repbase (http://www.girinst.org/repbase/) with bowtie (Langmead et al., 2009) using the options -M 1 -v 2. Transposons with fewer than 10 counts in each sample were excluded from further analysis. Counts were normalized to the total number of mapped reads for each library for the prg-1 dataset, or to the total number of non-ribosomal mapped reads for all other datasets. A pseudocount of 1 was added to each value to avoid division by zero errors. Analysis of sequencing data and plot creation was performed using the R statistical computing environment (R Core Team, 2013).

### Accession numbers

RNA-seq data reported in this study have been submitted to the GEO database and will be available at the time of publication.

### Immunofluorescence

Adult hermaphrodites raised at 20°C or 25°C were dissected in M9 buffer and flash frozen on dry ice before fixation for 1 min in methanol at −20°C. After washing in PBS supplemented with 0.1% Tween-20 (PBST), primary antibody diluted in in PBST was used to immunostain overnight at 4 °C in a humid chamber. Primaries used were 1:50 OIC1D4 (Developmental Studies Hybridoma Bank). Secondary antibody staining was performed by using a Cy3 donkey anti-mouse or Cy-5 donkey anti-rabbit overnight at 4°C. All images were obtained using a LSM 710 laser scanning confocal and were taken using same settings as control samples. Images processed using ImageJ. Marked P granule dysfunction was scored by eye as a loss of greater or less than 20% loss of P granules.

### DNA Damage Assay

Worms that were close to sterility were isolated and defined as sterile if they did not have any offspring as day 3 adults at 20°C or day 2 adults at 25°C. Fertile siblings of sterile worms were defined as ‘close to sterility’. The presence of the DNA damage response was determined by using a phospho-specific antibody targeting the phosphorylated consensus target site of ATM and ATR kinases (pS/TQ) (Cell Signaling Technology). This antibody has only been shown to stain a DNA damage response in the germline and was used as previously described (Vermezovic et al., 2012).

## Acknowledgments

We thank members of the Ahmed lab for critical reading of the manuscript and Jacinth Mitchell for wild type control, *nrde-1* and *nrde-2* RNA-seq data. Some strains were provided by the CGC, which is funded by NIH Office of Research Infrastructure Programs (P40 OD010440). This study was supported by NIH grants F32 GM120809 (K.B) and RO1 GM083048 (S.A). M.S. was supported by a DFG postdoc fellowship.

## Competing interests

Authors declare no competing interests.

## Author contributions

K.B., B.H., M.S. and S.F. performed experiments. S.F. analyzed the data. K.B., B.H., M.S., S.F. and S.A. wrote manuscript.

## Supplemental Information

**Fig. S1. Related to Figure 2.**
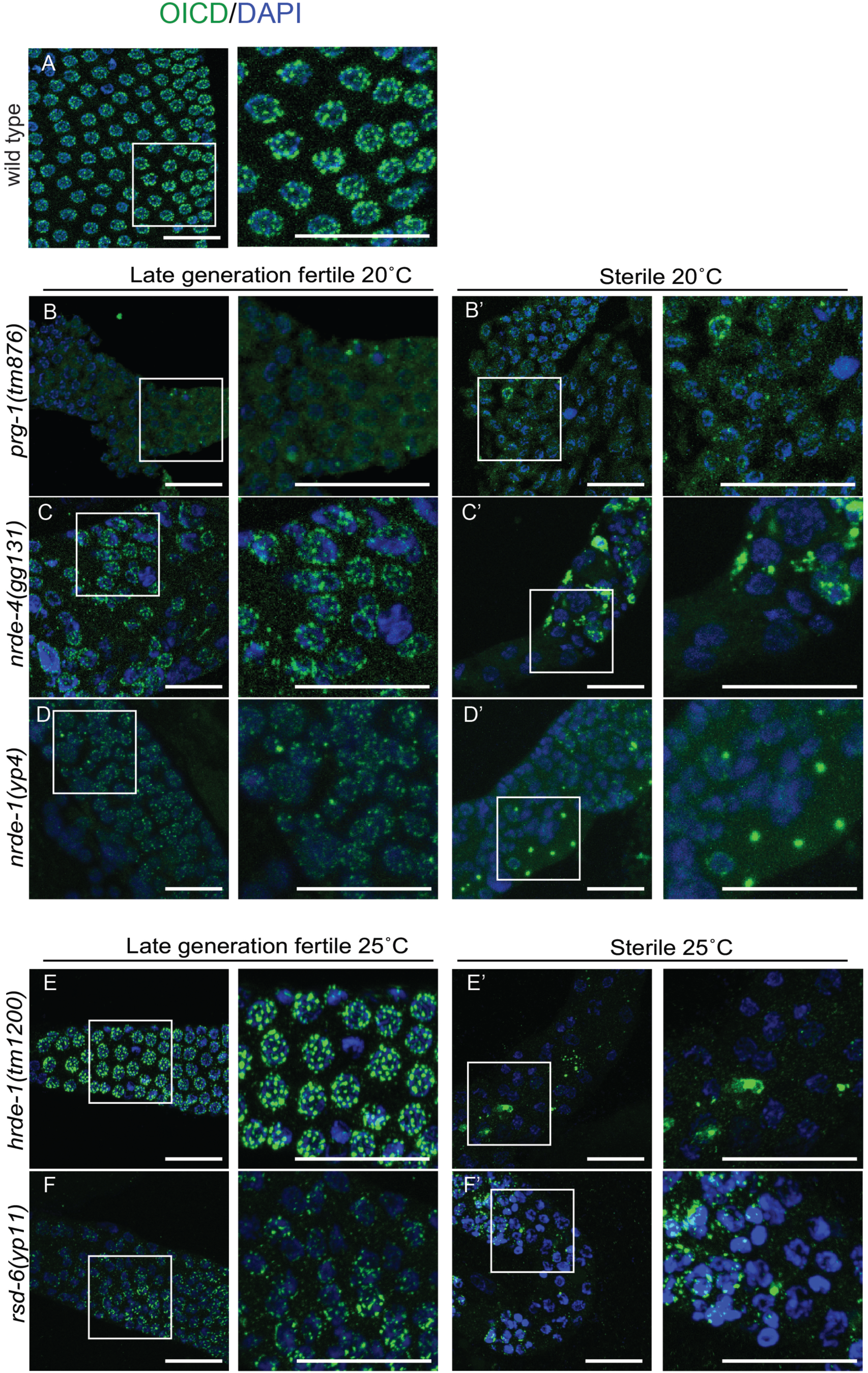
P granule defects in *prg-1, nrde-4, hrde-1 and nrde-1*. Germlines of sterile Day 2-3 adult animals were stained using the OIC1D4 antibody against P granules (green) and DAPI (blue). (A) Control animals contain uniform puncta of P granule staining surrounding each nuclei. (B-F) Late generation fertile *prg-1(tm876), nrde-4(gg131), nrde-1(yp4)*, *hrde-1(tm1200)* and *rsd-6(yp11)* and animals displayed P granule staining similar to wild type except for *prg-1(tm876).* (B’-F’) Sterile *prg-1(tm876), nrde-4(gg131), nrde-1(yp4)*, *hrde-1(tm1200)* and *rsd-6(yp11)* with P granule abnormalities.

**Table S1: Related to Figure 3.** Fertility of pgl-1 mutants at 25°C. 20°C P0 *pgl-1* mutants were shifted to 25°C and fertility was measured for the G1 generation. Fertile G1 worms were cloned and the fertility of the G2 generation was measured.

|  | Total Worms Scored | % Fertile | % Sterile |
| --- | --- | --- | --- |
| *pgl-1* G1 at 25°C | 91 | 22% | 78% |
| *pgl-1* G2 at 25°C | 71 | 0% | 100% |

**Table S2: Related to Figure 4.**
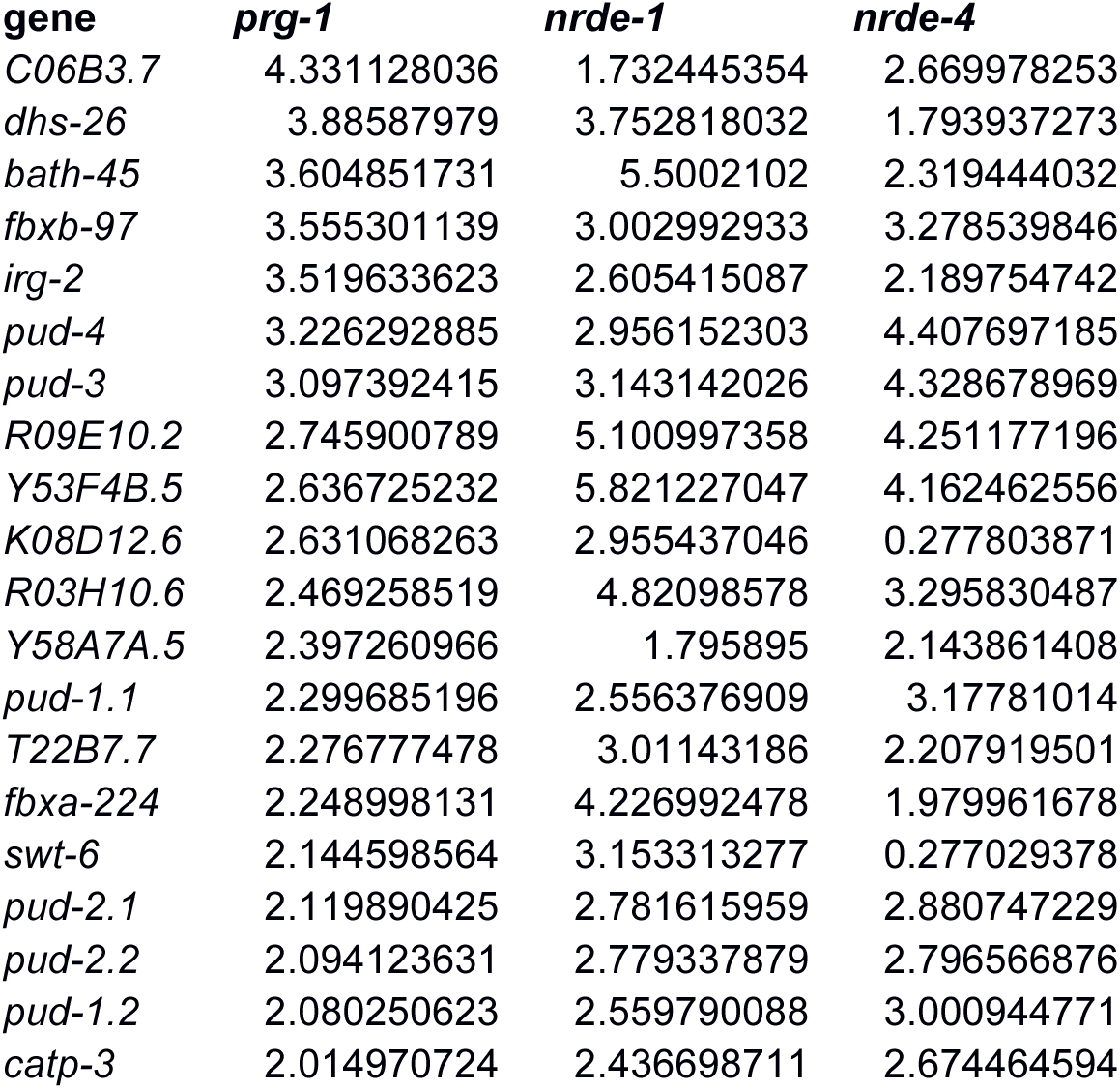
Genes upregulated in late-generation *prg-1* mutants and in sterile *nrde-1* and *nrde-4* mutant L4 larvae.

| gene | *prg-1* | *nrde-1* | *nrde-4* |
| --- | --- | --- | --- |
| *C06B3.7* | 4.331128036 | 1.732445354 | 2.669978253 |
| *dhs-26* | 3.88587979 | 3.752818032 | 1.793937273 |
| *bath-45* | 3.604851731 | 5.5002102 | 2.319444032 |
| *fbxb-97* | 3.555301139 | 3.002992933 | 3.278539846 |
| *irg-2* | 3.519633623 | 2.605415087 | 2.189754742 |
| *pud-4* | 3.226292885 | 2.956152303 | 4.407697185 |
| *pud-3* | 3.097392415 | 3.143142026 | 4.328678969 |
| *R09E10.2* | 2.745900789 | 5.100997358 | 4.251177196 |
| *Y53F4B.5* | 2.636725232 | 5.821227047 | 4.162462556 |
| *K08D12.6* | 2.631068263 | 2.955437046 | 0.277803871 |
| *R03H10.6* | 2.469258519 | 4.82098578 | 3.295830487 |
| *Y58A7A.5* | 2.397260966 | 1.795895 | 2.143861408 |
| *pud-1.1* | 2.299685196 | 2.556376909 | 3.17781014 |
| *T22B7.7* | 2.276777478 | 3.01143186 | 2.207919501 |
| *fbxa-224* | 2.248998131 | 4.226992478 | 1.979961678 |
| *swt-6* | 2.144598564 | 3.153313277 | 0.277029378 |
| *pud-2.1* | 2.119890425 | 2.781615959 | 2.880747229 |
| *pud-2.2* | 2.094123631 | 2.779337879 | 2.796566876 |
| *pud-1.2* | 2.080250623 | 2.559790088 | 3.000944771 |
| *catp-3* | 2.014970724 | 2.436698711 | 2.674464594 |

## Notes

#### Summary of Updates

Changes to the title, abstract, introduction, results and discussion were made in an effort to clarify the central messages of this manuscript.

